# Orion Microbiome Database: mapping constellations in the microbiome universe

**DOI:** 10.64898/2026.02.05.704118

**Authors:** Guangxi Wu, Yirui Tang, Grace Du, Nathan Heibeck, Julian Scaliogtti, William Francis, Shuiquan Tang

## Abstract

**Background:** Publicly available microbiome sequencing data have grown rapidly in scale and diversity, but secondary use remains limited by fragmentation across repositories, inconsistent annotations, heterogeneous processing pipelines, and incomplete metadata. These barriers complicate cross-study comparisons and restrict the reproducibility of large-scale microbiome research.

**Objective:** The Orion Microbiome Database was developed to overcome these challenges by providing a standardized, accessible, and reproducible resource for the integration and analysis of whole-genome shotgun (WGS) microbiome datasets.

**Methods:** Orion aggregates raw metagenomic datasets from public repositories and harmonizes them into computed microbial profiles using automated and transparent analysis pipelines. Metadata are curated to ensure consistency and enable systematic cross-cohort comparisons. The platform provides interactive tools for data exploration, visualization, and downstream analysis, reducing redundancies in data preprocessing and lowering the barrier for both novice and expert users. Beyond research applications, Orion also serves as an educational resource, enabling microbiome data exploration in classroom settings.

**Conclusion:** By unifying fragmented sequencing resources into a reproducible framework, Orion accelerates microbiome discovery, supports scalable cross-study investigations, and fosters integration of microbiome science into teaching and training. This database has the potential to serve as a model for future large-scale, community-driven platforms in microbiome research.

## Introduction

Microbiome databases are crucial in enabling large-scale data integration, advancing our understanding of host-microbiome interactions, and facilitating translational discoveries. Thus, we propose the Orion Microbiome Database, a comprehensive resource designed to consolidate and standardize microbiome data for enhanced accessibility and analysis. In its first iteration, we focus on the gut microbiome, arguably the most well-studied microbial community, given its pivotal role in host physiology, including immune function (1,2), metabolism (3,4), energy production (5,6), and disease susceptibility (7–14).

The gut microbiome, unlike many other microbiomes, is shaped by diverse factors, including age (15), sex (16), BMI (17), geography (18), genetics (19), diet (20), and antibiotic use (21), necessitating robust databases to unravel these complex relationships (22). With the rapid expansion of gut microbiome research and other microbiome research, large-scale metagenomic datasets—including 16S amplicon and shotgun sequencing data—have been generated and deposited in public repositories such as the European Nucleotide Archive (23), NCBI Sequence Read Archive (SRA) (24), and processed databases like MGnify (25), gcMeta (26), Qiita (27), and NMDC (28) Additionally, specialized resources such as gutMDisorder (29), GIMICA (30), GMrepo (31,32), MicrobiomeDB (33), and DISBIOME (34) link microbial dysbiosis to diseases, while curatedMetagenomicData (35) offers standardized taxonomic and functional annotations.

Despite these advances, significant challenges remain in data accessibility and reusability within this field. These challenges include: incomplete or inconsistent metadata—such as missing age, sex, or BMI—which hampers cross-study comparisons; the lack of intuitive, interactive tools for real-time data exploration and the lack of integrated analysis pipeline, limiting their utility for researchers without advanced computational skills; the lack of dynamic visualization or collaborative features that enhance user engagement and data interpretation; the inability to submit customer data to be analysed alongside data in the repository. All the aforementioned existing databases suffer from one or more of these shortcomings.

To address these limitations, **Orion** combines expert-curated shotgun sequencing data with community-driven innovation to improve accessibility, interactivity, and collaboration in microbiome research. Initially developed through extensive manual curation by scientists at Zymo Research, Orion ensures high-quality, standardized metadata and analysis by applying a single standardized analysis pipeline to enhance reproducibility. Unlike static repositories, Orion allows users worldwide to contribute open-source data, build custom cohorts, and collaborate on shared projects, fostering a continuously expanding knowledge base. These unique features help address critical gaps in current microbiome resources.

In addition to its community-driven data integration, Orion provides a sophisticated, user-friendly interface for dynamic data filtering, summary visualization, and feature selection, enabling researchers to efficiently explore large metagenomic datasets. The platform also supports interactive taxonomic composition analysis at all taxonomic levels (from Kingdom to Species) across three major domains (Eukaryotes, Prokaryotes, Viruses), with customizable visualizations that update in real time based on user-selected data points. Recognizing the importance of shared scientific discovery, Orion integrates features that facilitate seamless collaboration, allowing users to share, annotate, and discuss datasets within the platform. By combining expert-curated data with crowd-sourced contributions, intuitive visualization, and collaborative capabilities, Orion aims to democratize microbiome research and accelerate the curation process, making complex metagenomic data more accessible, interpretable, and actionable for researchers worldwide.

## Methods

### Data Discovery, ingestion and curation

A flowchart of data discovery, ingestion and curation is demonstrated in **Figure 1**. Data was identified and collected from several different public repositories but was primarily accessed through NCBI using references to BioProject IDs (https://www.ncbi.nlm.nih.gov/bioproject). A list of potentially valid BioProject IDs was created through a team effort of extensive web searches for scientific publications using keywords relevant to the database criteria. Primary keywords in the search were for “fecal”, “gut”, or “stool” samples, a phenotypic pattern such as “Parkinsons” or “irritable bowel syndrome”, and a data type such as “shotgun” or “WGS”. Identified publications were screened for the presence of a BioProject ID linked to National Center for Biotechnology Information (NCBI) or the European Nucleotide Archive (ENA) (https://www.ebi.ac.uk/ena/browser/home). We also referenced existing data repositories for the gut microbiome such as GMrepo (31,32) and gutMEGA (36), cross referencing BioProject IDs present in their catalogue with our key word discovery list, adding any unique IDs not yet catalogued.

**Figure 1.**
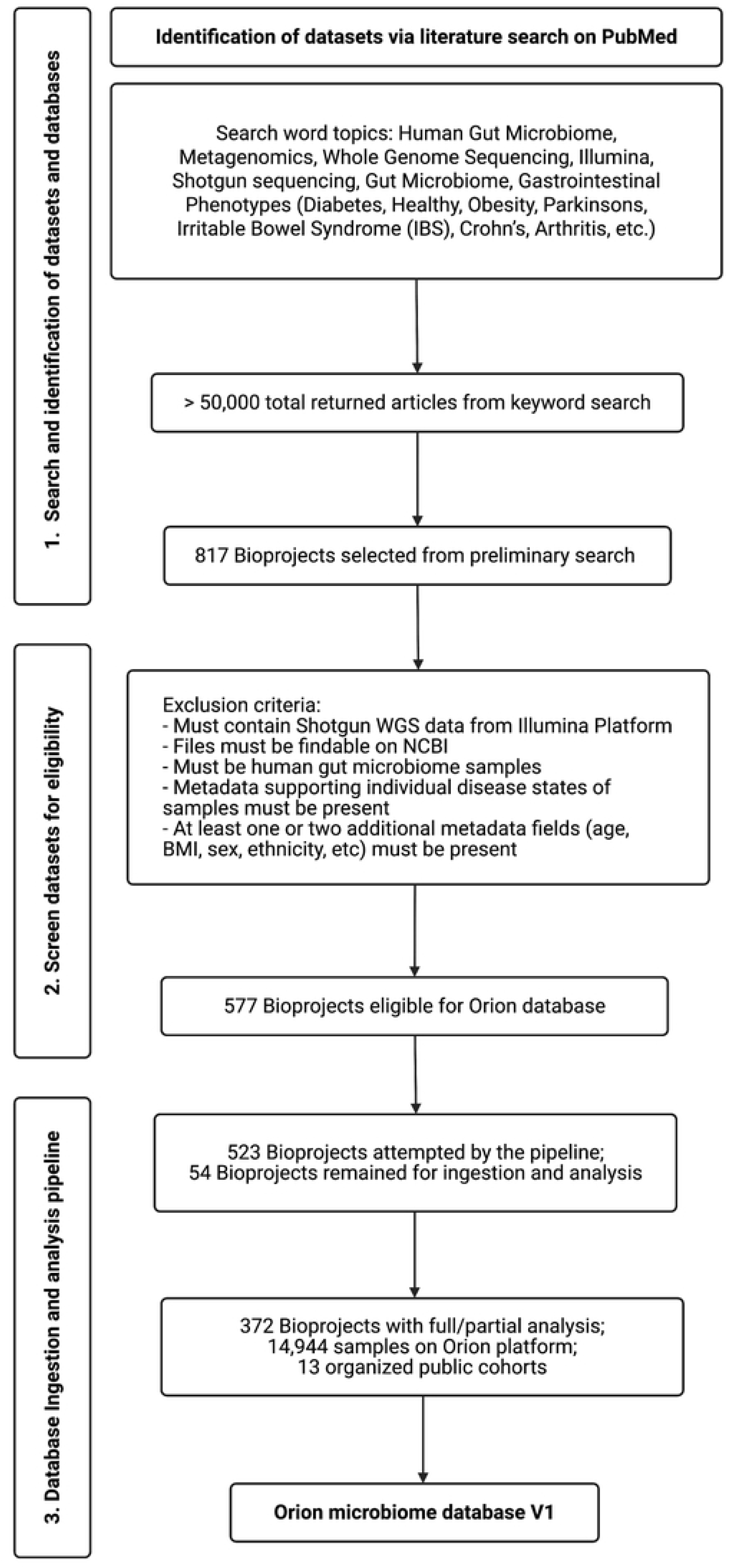
Flowchart of data discovery, ingestion and curation for Orion Microbiome Database V1.

After cataloguing a BioProject ID correlated to a publication, the articles were separately screened for impact and validity of data ingestion. Screening criteria included proper sequencing methods prioritizing Illumina Whole Genome Sequencing (WGS), complete metadata with per-sample classifications for harmonization, and a clearly defined methodology of participant selection and sample collection. BioProject IDs within papers that satisfied these criteria were then fed into an internal workflow that requests all associated information from NCBI or ENA.

Sample uploads to either repository was not always congruent with the publication due to mixed metadata, unidentified samples, missing samples, or alternative sequencing methods sometimes being present following the fetching process. Immediately following any successful fetch, the acquired metadata is examined to account for proper sample type, quantity of samples, and sample identification. Non-WGS samples and any unidentifiable samples are removed.

NCBI metadata often incorrectly identifies sample group for some samples and is often missing metadata fields that may be present in the original publication. Following sample QC, all present samples are harmonized by filling in missing fields if available in published documentation and normalizing existing fields in terminology to adhere to an agreed upon structure and nomenclature. Orion prioritizes gathering as much metadata as possible and categorizes metadata into two groups. Primary metadata are the most common fields documented consistently across studies. These fields exist in a structured format within Orion, making them directly able to be queried directly by the user and normalized for direct comparisons. Secondary metadata fields are any fields present within the NCBI/ENA fetch that do not match one of the designated primary metadata fields. These fields and the entries within them are preserved, but due to the sparse and inconsistent nature of this information across studies, they are not normalized or stored in a structured way to be immediately queried later. A list of primary and secondary metadata fields and the terminology/methodology used to standardize each can be found in **Table 1**.

**Table 1.**
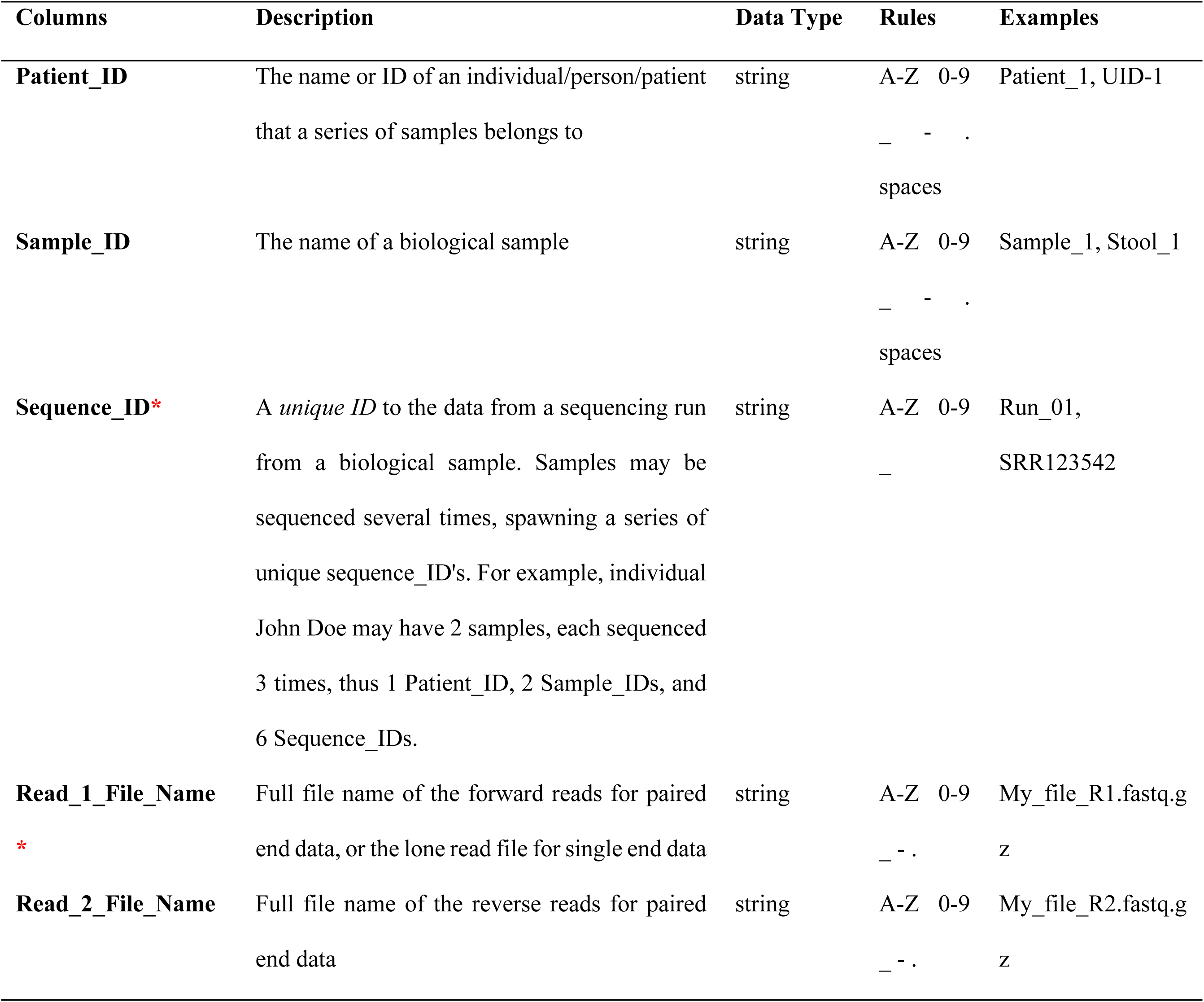
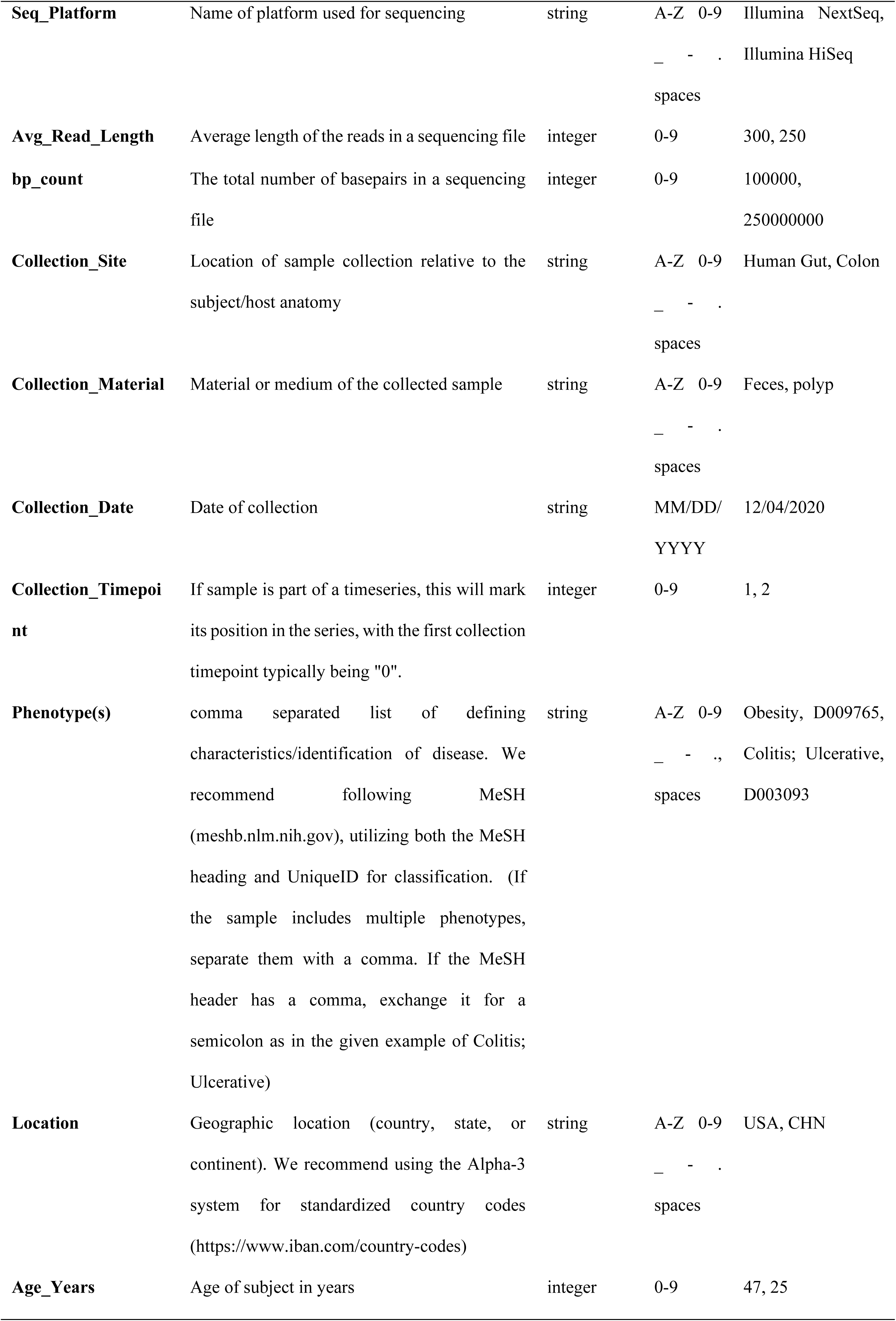
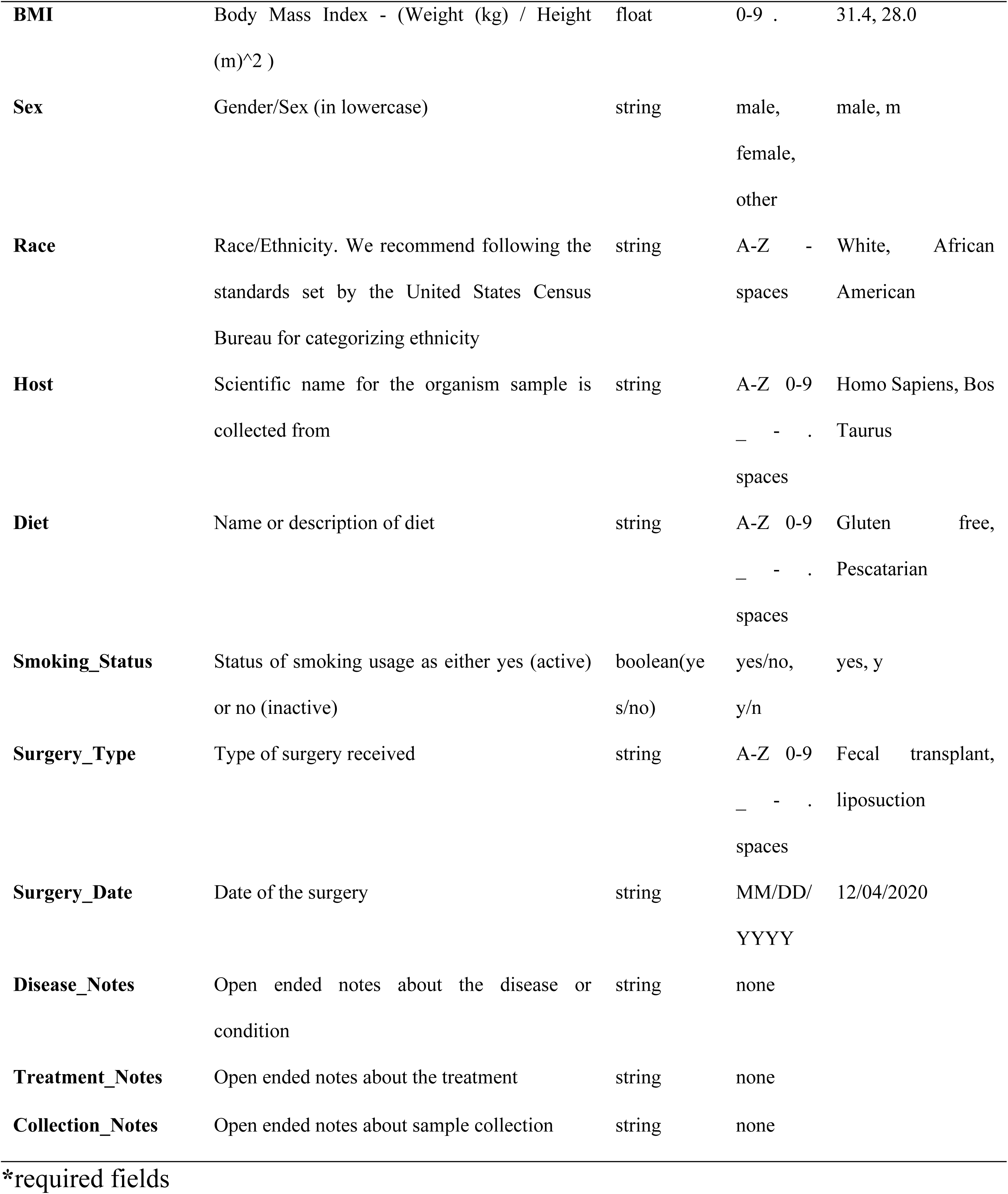
Orion Metadata Template and Guide.

An ingestion template based on the criteria stated in **Table 1** is provided to all users on Orion. Orion encourages but does not enforce strict adherence to these guidelines for private data, but all public data curated internally by the Orion team or submitted for public use by private users is reviewed to ensure adherence to this format to preserve downstream comparability among individual samples.

Following the completion of metadata harmonization, samples are submitted to an internal standardized pipeline for processing. All curation efforts were focused on publications prior to June 2025. Ongoing data curation and ingestion of additional data for gut as well as other human sampling sites is planned for future iterations of Orion.

### Bioinformatics Pipeline

All taxonomic analysis on Orion is standardized using a single analysis pipeline performed using Zymo’s internal High-Powered-Compute (HPC) cluster. The bioinformatics pipeline used to conduct taxonomic profiling was adapted from nf-core/taxprofiler version 1.0.0 (37) The most significant change relevant to this study is the use of sourmash v4.8.2 as the default taxonomy profiling tool (38). A full list of changes and source code can be found at https://github.com/Zymo-Research/aladdin-shotgun. All data in Orion database are processed with this pipeline using the following steps and parameters:

The reads were adapter- and quality-trimmed using fastp 0.23.4 (39). Reads shorter than 15bp were discarded after trimming. Low complexity reads were removed using BBDuk 39.01 (https://sourceforge.net/projects/bbmap/) with the parameter ‘entropy=0.3 entropywindow=50’. Kmer sketches of each sample were then produced with sourmash v4.8.2 with the parameter ‘k=31,scaled=1000,abund’. The sketches were then searched against a sourmash database mainly consisting of GTDB R08-RS214 genomic representatives (40), Genbank viral, archaeal, protozoa, fungi genomes, and common host genomes, with the parameter ‘--ksize 31 --threshold-bp 5000’. The detailed list and data download source can be found at https://zymo-research.github.io/pipeline-resources/about_pages/about_shotgun.html. We estimate the numbers of reads belonging to each microbial genome by multiplying ‘f_unique_weighted’ values of that genome from sourmash to the total number of reads input to the sourmash step. Reads assigned to host genomes were removed from downstream analysis. Samples with less than 1000 reads assigned to microbial genomes were excluded. The read counts of each microbial genomes for each sample were used as input for constructing sample profiles on Orion database.

### Cohort building

Cohort creation is an integral part of the Orion platform allowing researchers to directly build upon previous curation work. Long-termly, we invite active users to publish their private curation work for others to utilize. Internally, our curation team assembled 13 preliminary public cohorts for various diseases, demographics, and health states comprised of samples from the public domain. Cohort curation followed a general workflow of identifying a specific phenotype, such as a disease state, using the MeSHID label (https://meshb.nlm.nih.gov/) and selecting any matching samples. Then, further curation using associated metadata can restrict the subgroup down to specific age, sex, geographical region, etc, which is documented in the cohort summary details hosted on Orion. A summary of the 12 public cohorts can be explored in **Supplementary Table 1.**

### Statistical analysis

The Orion platform integrates pre-computed taxonomic profiles to enable downstream statistical and exploratory analyses through a suite of plug-in modules. Current functionality includes taxonomy-based queries, taxonomic composition bar plots across seven hierarchical levels, and three-dimensional principal coordinates analysis (PCoA) plots of beta diversity. Additional features under development include differential abundance testing, alpha diversity estimation, and biomarker discovery, which are undergoing internal validation prior to release.

Several modules are currently available on Orion. Taxonomic bar plots are generated through python3 and visualized through Nivo (41,42). Interactive filtering options allow users to select specific taxonomic levels or subsets of taxa for visualization. PCoA plots are computed dynamically for user-defined cohorts, with Bray–Curtis dissimilarity used to construct distance matrices and ordinations generated via Scikit-bio (43). Taxonomy-based queries are executed against Orion’s internal reference database, enabling sample retrieval based on the presence, absence, or relative abundance of specified taxa to facilitate targeted cohort construction.

Planned modules for differential abundance and biomarker analysis will employ LEfSe with standard thresholds (p ≤ 0.05 and linear discriminant analysis [LDA] score ≥ 2) for biomarker detection and visualization, including cladogram summaries (44). We applied these tests in our case studies. Additional differential abundance testing and multivariate association analyses will be conducted using MaAsLin3, applying default parameters for data normalization, false discovery rate (FDR) control, and abundance/prevalence thresholds (45). Alpha diversity metrics will be estimated using Qiime2, with support for three commonly used indices: Shannon, Chao1, and Simpson (46). All analysis modules within Orion are implemented using open-source software with standardized, transparent parameterization to ensure reproducibility.

### Database construction and web development

All data were loaded into a PostgreSQL database, interfaced, accessed, and queried through python3 (47,48). The frontend (webpages) of the website was coded using React (49) and a combination of typescript/javascript/html tools, while the backend was coded using a python3 FastAPI (RESTful), celery asynchronous task manager, and AWS lambda functions (50–52). Plotly was used for 3D visualizations on the front-end (53). Various other open-source JavaScript libraries were also used in front end utility and visualization. The website is hosted on AWS using ECS (54).

## Results

### Data Processing

In totality, our searches yielded 814 potential BioProjects/publications to examine. From the initial 814 BioProjects identified, 588 met the criteria for attempted fetching, while 28 did not. An additional 198 remain unexamined, not yet evaluated by the curation team. Of the 588 BioProjects fetched, 216 failed initial QC and were omitted from further processing, leaving 372 BioProjects successfully ingested for further processing. A total of 23,521 samples from 166 of the 372 BioProject were ingested, harmonized, and submitted to the pipeline on Orion. Of these, 14,824 samples successfully completed analysis, with the other 8,679 failing to produce pipeline results for various reasons. The remaining 206 BioProject that are ingested but not analysed are scheduled to be fed through the pipeline as additional compute and personnel resources become available.

### Case study 1: Reproducibility of Colorectal cancer vs Healthy Adults results from PRJNA447983

In case study 1, a colorectal cancer (CRC) dataset was selected to evaluate Orion’s fidelity during data integration through reproducing the analyses performed in the initial study (55). This paper was a meta-analysis study of fecal metagenomes across several CRC cohorts, including BioProject PRJNA447983 which was the dataset uploaded to Orion. This upload comprised of two cohorts: cohort 1 containing 29 CRC and 24 healthy samples, and cohort 2 containing 32 CRC and 28 healthy samples. A summary of the subject characteristics is provided in **Table 2**.

**Table 2.** CRC sample characteristics.

| Dataset | Groups (N) | Age (year, mean ± s.d.) | BMI (mean ± s.d.) | Sex F(%) / M(%) | Country | # of reads (10E9) |
| --- | --- | --- | --- | --- | --- | --- |
| Cohort 1 | Control (24) | 67.9 ± 7.1 | 25.3 ± 3.5 | 45.8/54.1 | Italy | 8.2 |
|  | CRC (29) | 71.4 ± 8.2 | 25.7 ± 4.1 | 20.7/79.3 |  |  |
| Cohort 2 | Control (28) | 57.8 ± 8.3 | 24.6 ± 3.8 | 42.9/57.1 | Italy | 5.1 |
|  | CRC (32) | 58.4 ± 8.4 | 26.8 ± 4.3 | 28.1/71.9 |  |  |

Species richness and differential abundance analyses were performed to compare with results from the original paper. The original study reported species richness of cohort 1 to have a median of 80 species observed and interquartile range (IQR) from 67 to 87 for healthy individuals and a median of 81 and IQR from 62 to 88 for individuals with CRC (p-value = 0.922, assessed by two-tailed Wilcoxon rank-sum tests). For cohort 2, the original study reported a median of 99, a IQR from 92 to 110 for healthy individuals and a median of 120 and IQR from 110 to 130 for individuals with CRC (p-value = 0.001). The Orion reanalysis results of these samples report for cohort 1 a median species richness of 225 and IQR from 190 to 325 for healthy individuals, and a median of 195 and IQR from 115 to 298 for individuals with CRC (p-value = 0.216). For cohort 2 a median species richness of 240 and IQR from 190 to 405 for healthy individuals, and a median of 460 and IQR from 360 to 510 for individuals with CRC (p-value = 0.001). These outcomes align with the original study’s results, which also reported no significant difference in species richness between cohort 1 healthy and CRC groups (p-value = 0.922), while finding significant difference between cohort 2 healthy and CRC groups (p-value = 0.001), demonstrating a level of reproducibility of the cohort-specific diversity pattern when compared to the original publication, despite differences in absolute richness values between the two analyses (**Figure 2**).

**Figure 2.**
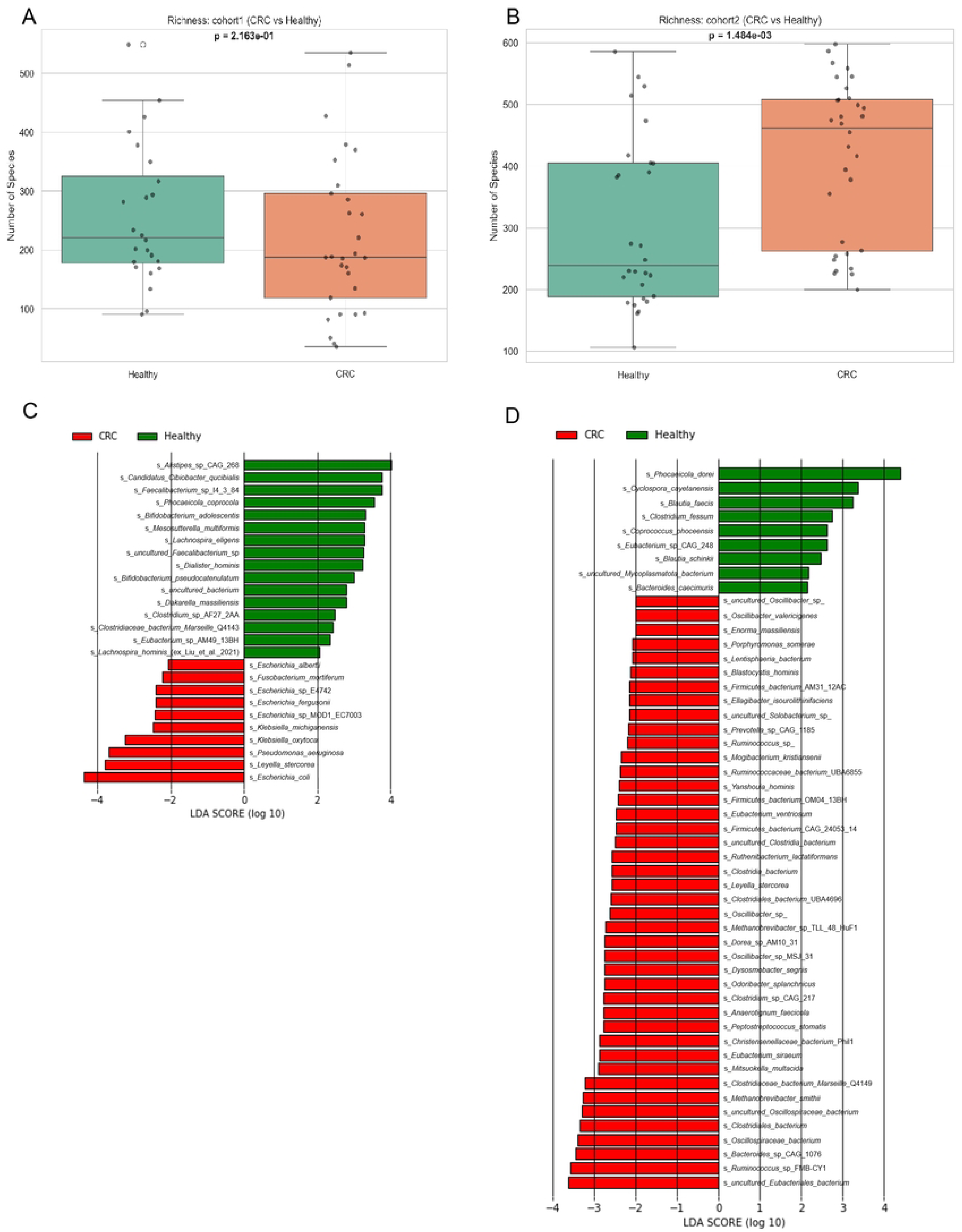
Boxplots of species richness and LDA plots of relative abundances generated from Orion dataset. (A) Boxplot for cohort 1 showing species richness. P-values are calculated by two-tailed Wilcoxon rank-sum tests; (B) boxplot for cohort 2; (C) relative abundance and effect sizes estimated using LDA score in LEfSe for the significantly different microbial species in CRC samples compared to healthy samples for cohort 1; (D) for cohort 2.

Next, we examined differential relative abundances of microbial species in CRC versus healthy samples using LEfSe, which is the same tool used in the original paper (55). Both analyses identified ten significantly enriched organisms in CRC when compared to healthy with four common species, *Escherichia coli*, *Leyella stercorea*, *Klebsiella oxytoca*, and *Fusobacterium mortiferum*, of which the first two are among the top three most enriched organisms in both analyses (**Figure 2C**). In Cohort 2, the original study reported 19 significantly enriched taxa in CRC, whereas the Orion analysis identified 42. Among these, four common species were found: *Eubacterium ventriosum, Odoribacter splanchnicus, Peptostreptococcus stomatis,* and *Methanobrevibacter smithii*, which covers four out of the top six identified in the original study (**Figure 2D**). With the goal being to assess reproducibility between this LEfSe analysis and the original study, these findings suggest partial reproducibility rather than full concordance.

### Case study 2: Parkinson’s disease cohort

In case study 2, we focus on Parkinson’s disease (PD) phenotype. In total, 890 samples on Orion had PD as the phenotype; further, 185 samples were chosen from age-matched healthy individuals. A summary of the subject characteristics is provided in **Table 3**. These samples come from three BioProjects, each using a different sequencing platform (Illumina NovaSeq 6000, Illumina HiSeq 2500, and Illumina HiSeq X Ten). While 179 samples from the project were from female subjects, 309 samples were from male subjects. The average age of the samples was 68.67 years, with an average BMI of 27.96 kg/m^2^.

**Table 3.** Sample characteristics of case study 2.

| Characteristics | Data in Summary | Data by Each BioProject |  |  |  |  |  |  |  |
| --- | --- | --- | --- | --- | --- | --- | --- | --- | --- |
|  |  | PRJEB17784 |  | PRJNA834801 | PRJNA743718 | PRJNA433459 |  | PRJEB53401 | PRJEB53403 |
|  |  | Parkinson's Disease | Healthy | Parkinson's Disease | Parkinson's Disease | Parkinson's Disease | Healthy | Parkinson's Disease | Parkinson's Disease |
| No. of samples | 1075 | 216 | 145 | 488 | 82 | 38 | 40 | 48 | 18 |
| Avg age (year) | NA | NA |  | 68.67 ± 8.54* |  | 66.6 ± 7.1* |  |  |  |
| Sex |  |  |  |  |  |  |  |  |  |
| Female |  | 0 | 0 | 179 | NA | 21 | NA | 22 | NA |
| Male |  | 216 | 145 | 309 | NA | 17 | NA | 26 | NA |
| BMI (kg/m<sup>2</sup>) | NA | NA |  | 27.96 ± 5.40* |  | NA |  |  |  |
| Sequencing platform |  | Illumina HiSeq 2500 | Illumina HiSeq 2500 | Illumina NovaSeq 6000 | Illumina NovaSeq 6000 | Illumina HiSeq X-ten | Illumina HiSeq X-ten | Illumina NextSeq 550 | Illumina NovaSeq 6000 |

\*Values are means ± SD

To investigate the potential associations between the fecal microbial taxonomic abundance and different sequencing platforms in subjects with PD, multivariate generalized linear models using MaAsLin3 were performed on the differential taxa (minimal abundance = 0.01, minimal prevalence = 0.20) and sequencing platforms. The PRJNA834801 project with Illumina NovaSeq 6000 had the highest relative abundances of species *Cyclospora cayetanensis, Blautia wexlerae,* and *Blautia faecis*, and the lowest relative abundance of species *Alistipes putredinis* (FDR < 0.05 for all) (**Figure 3**). The project PRJNA433459 using Illumina HiSeq X-ten had the highest relative abundances of species *Bacteroides thetaiotamicron*, *Bacteroides ovatus*, *Bacteroides uniformis,* and *Phocaeicola dorei* and the lowest abundances of *Akkermansia muciniphila*, *C. cayetanensis*, *Fusicatenibacter saccharivorans*, *Candidatus Cibiobacter qucibialis*, uncultured *Eubacteriales* bacterium, *B. wexlerae* and *B. faecis* (FDR < 0.05 for all) (**Figure 3**).

**Figure 3.**
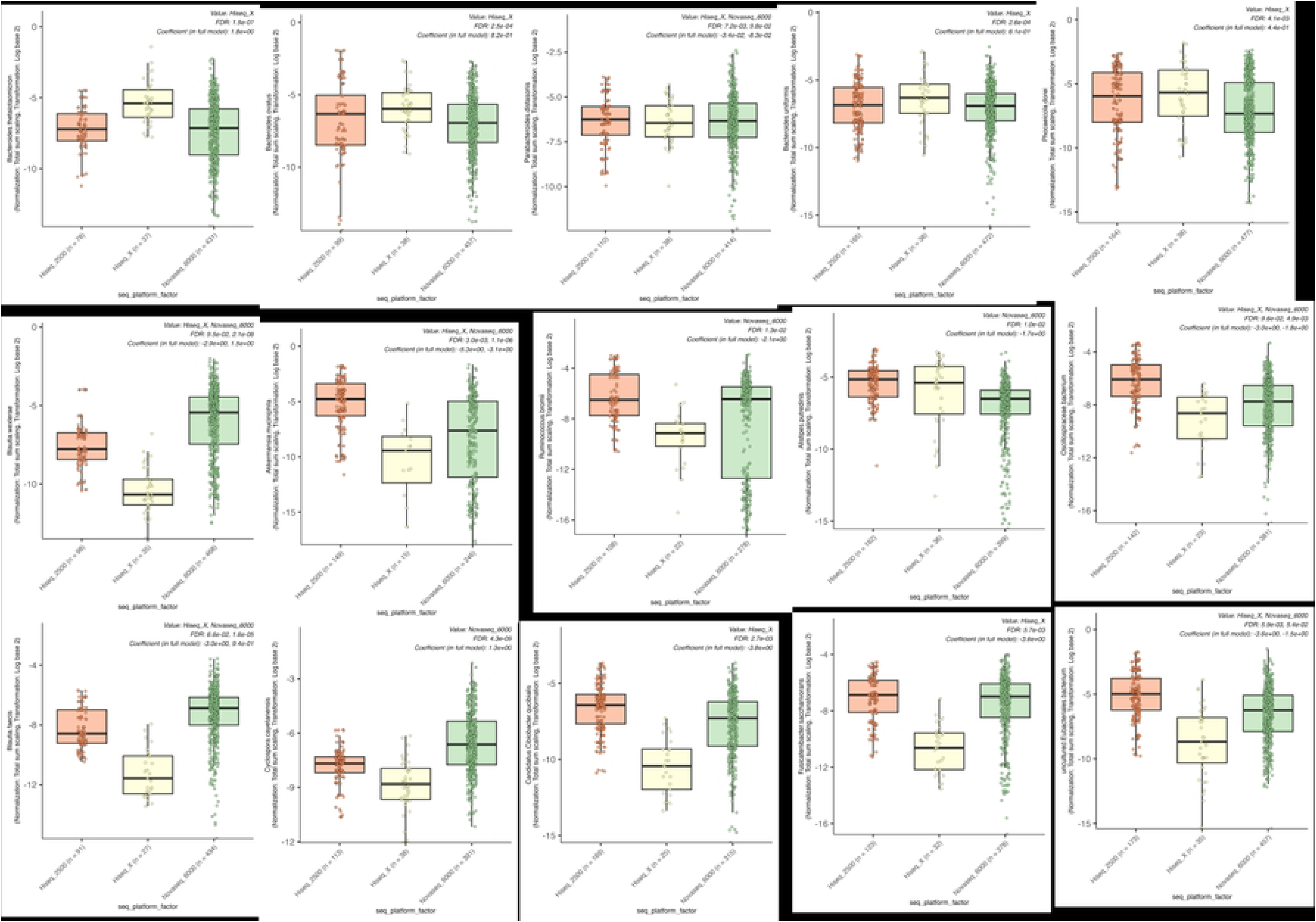
Boxplots of species with significant differences by sequencing platforms in the Parkinson disease cohort.

We then investigated the differences in the fecal microbial taxonomic abundance between subjects with PD and healthy phenotype using the multivariate generalized linear models in MaAsLin3 (minimal abundance = 0.01, minimal prevalence = 0.20). Comparing to healthy subjects, individuals with PD had significantly higher relative abundances of *B. wexlerae, C. cayetanensis, Clostridiales* bacterium, uncultured *Oscillospiraceae* bacterium, uncultured *Eubacteriales* bacterium, and *F. saccharivorans* (FDR < 0.05 for all) as well as lower relative abundances of *B. thetaiotamicron* and *Phocaeicola massiliensis* (FDR < 0.05 for all) (**Figure 4**).

**Figure 4.**
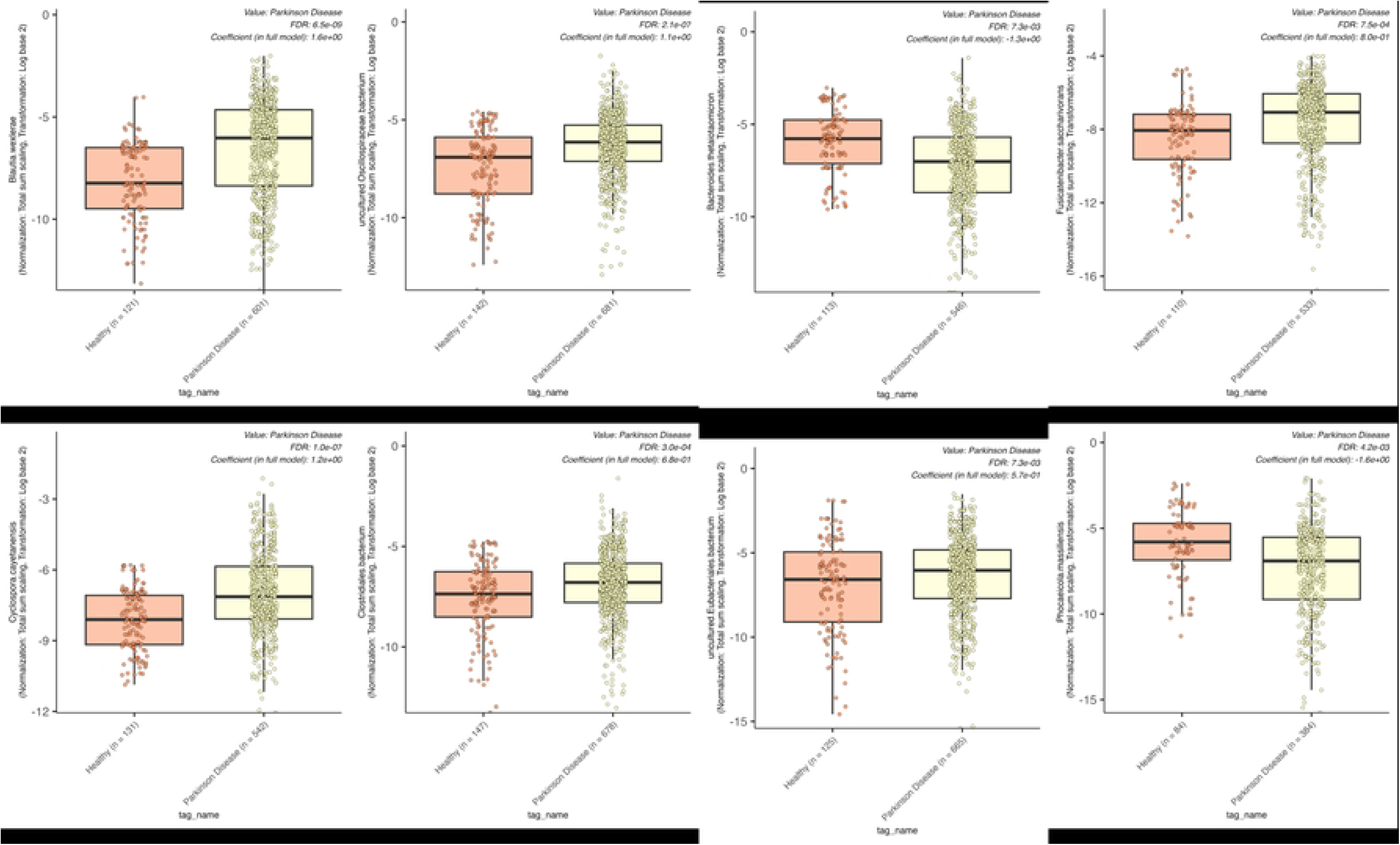
Boxplots of species with significant differences between healthy and Parkinson’s disease cohorts.

### Case Study 3: Colorectal cancer prediction model

Colorectal cancer (CRC) represents a significant global health burden, with delayed diagnosis often resulting in poor clinical outcomes (56,57). Early detection through non-invasive screening methods could substantially improve survival rates (58). Traditional screening approaches like colonoscopy, while effective, suffer from limited patient compliance due to their invasive nature (59,60).

This case study aimed to develop machine learning models capable of distinguishing between healthy individuals, patients with adenomas (precancerous polyps), and those with colorectal cancer based on gut microbiome taxonomic profiles. By explicitly modeling the adenoma stage—a critical intervention point in CRC progression—we sought to create clinically relevant risk stratification tools that capture disease progression beyond traditional binary classification approaches.

Using Orion’s cohort-building functionality, we identified and extracted data from participants across multiple studies with clearly defined phenotypic classifications. Two cohorts were curated using standardized filters: age ≥ 30 years, shotgun sequencing depth >1M reads, complete metadata, and diverse geographical origins. This yielded 2,392 samples comprising 1,705 healthy controls, 198 adenoma cases, and 489 CRC patients (**Figure 5**).

**Figure 5.**
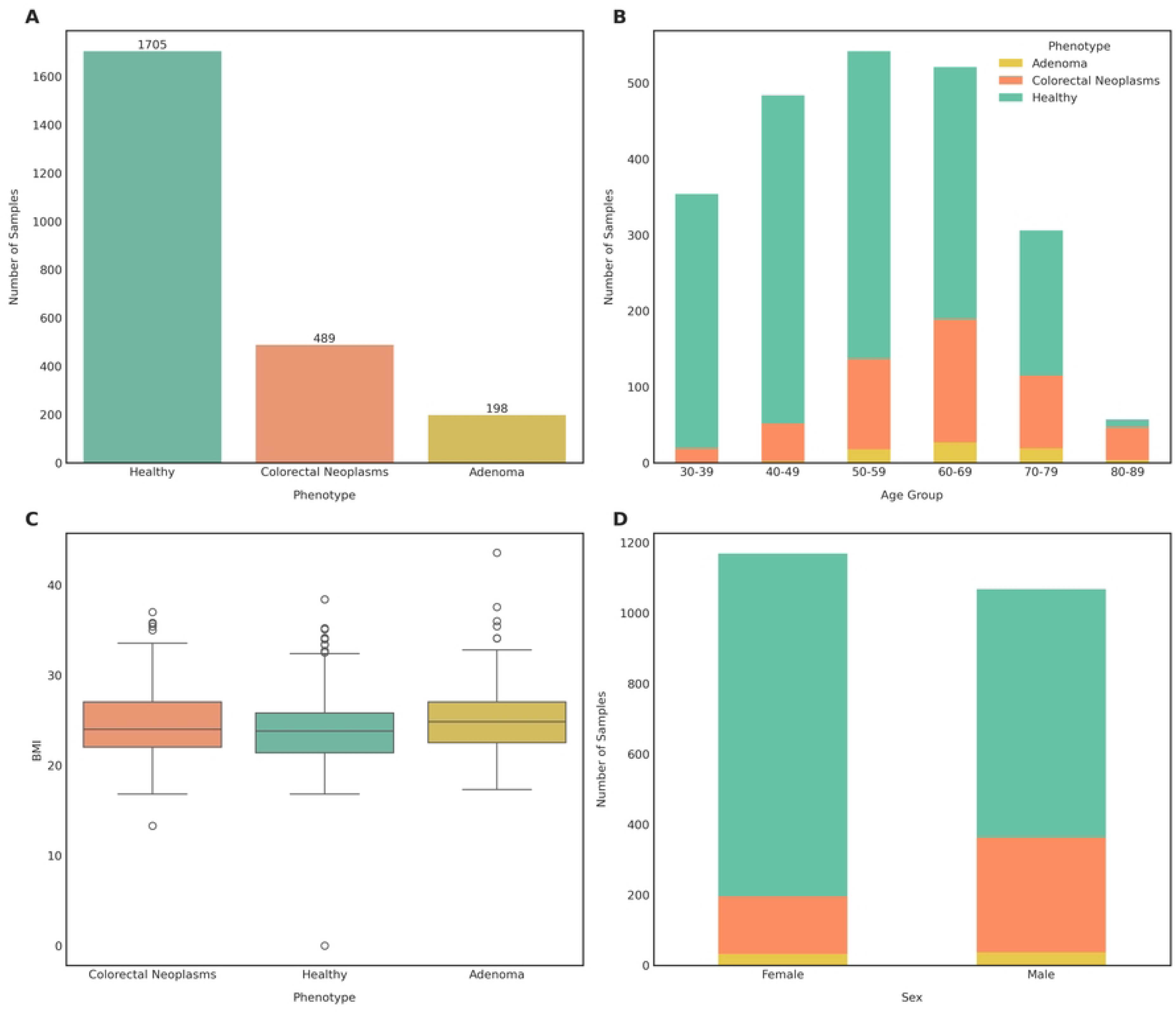
Sample composition and demographic characteristics of the microbiome cohort. **(A)** Sample distribution by phenotype. **(B)** Age distribution by phenotype. **(C)** BMI distribution by phenotype. **(D)** Sex distribution by phenotype.

All taxonomic profiles underwent standardized preprocessing to ensure data quality and biological relevance. Centered log-ratio (CLR) transformation addressed the compositional nature of microbiome data. Low-prevalence taxa (present in <5% of samples) were filtered to reduce noise. Non-biological confounders, including geographical coordinates, collection dates, sex, and sequencing metadata, were excluded to prevent spurious associations. Remaining numerical features were z-score standardized, and categorical variables were one-hot encoded.

We implemented two complementary modeling approaches. The binary classification framework distinguished healthy individuals from CRC patients using healthy and CRC samples. The multiclass framework extended this by incorporating the adenoma stage, classifying samples into three categories (healthy, adenoma, or CRC) using the complete 2,392-sample dataset. Both approaches employed three algorithms: logistic regression (with L2 regularization for binary; multinomial formulation for multiclass), random forest (600 trees, balanced subsample weights), and histogram gradient boosting (adaptive learning rate, no L2 regularization). Class imbalance was addressed through balanced class weighting. Model performance was evaluated using five-fold stratified cross-validation, with the best-performing algorithm subsequently validated on a stratified 25% held-out test set. All analyses used a fixed random seed to ensure reproducibility.

Both modeling approaches incorporated probability-based risk stratification systems tailored to their classification schemes. The binary model categorized individuals into four risk levels (low, medium, high, and CRC) based on predicted CRC probability thresholds (0.15, 0.35, and 0.50). The multiclass model employed a classification-based system that assigns risk categories based on the predicted class and model confidence: samples predicted as CRC are classified as High Risk, those predicted as adenoma are classified as High Risk, and those predicted as healthy are classified as Medium or Low Risk depending on prediction confidence.

The binary classifier achieved an AUROC of 0.960 in cross-validation and 0.947 on the held-out test set, demonstrating consistent discrimination across both evaluation settings (**Table 4**). The multiclass model achieved overall accuracy of 84.4% and effectively distinguished all three phenotypic states. Adenoma detection achieved 94.1% recall with 87.5% precision, while healthy and CRC classes maintained balanced precision-recall profiles (**Table 5**). All three classes achieved AUROCs exceeding 0.91 (**Table 5**).

**Table 4.** Performance metrics of the binary model using the histogram gradient boosting algorithm on the held-out test set and five-fold cross-validation.

| Metric | Held-out Test Set | Cross-validation |
| --- | --- | --- |
| Accuracy | 88.5% | 90.0% |
| Balanced Accuracy | 79.2% | 82.1% |
| Precision (CRC) | 81.7% | 84.7% |
| Recall (CRC) | 62.3% | 67.9% |
| F1-score (CRC) | 0.707 | 0.751 |
| AUROC | 0.947 | 0.960 |

**Table 5.** Performance metrics of the multiclass model by class.

| Class | Precision | Recall | F1-score | AUROC |
| --- | --- | --- | --- | --- |
| Healthy | 72.2% | 65.5% | 0.687 | 0.924 |
| Adenoma | 87.5% | 94.1% | 0.907 | 0.926 |
| CRC | 60.0% | 11.5% | 0.194 | 0.917 |

*Note: Probability re-weighting (2x weight on the CRC class) improved CRC recall from 11.5% to 65.5% at the cost of reduced adenoma recall (94.1% to 11.5%), demonstrating the trade-off inherent in addressing severe class imbalance. AUROC values remain unchanged as they measure ranking ability independent of classification thresholds.*

Risk stratification analysis demonstrated clinically meaningful separation of phenotypes (**Figure 6A**). The High Risk category was predominantly composed of CRC patients (69.9%), with 66.4% of all CRC samples correctly classified in this risk tier. In contrast, the Medium Risk category consisted primarily of healthy individuals (87.5%), reflecting lower disease probability predictions.

**Figure 6.**
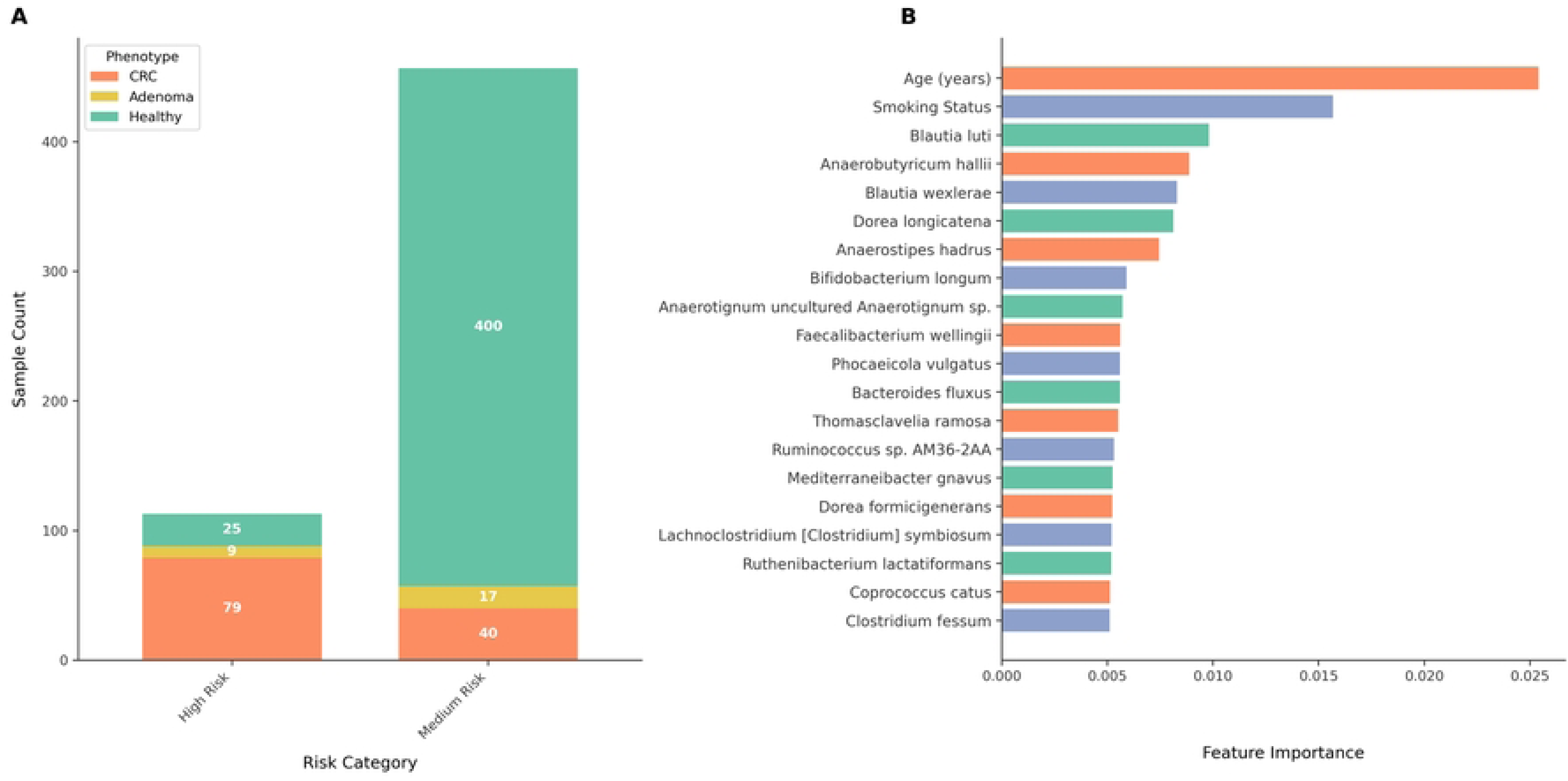
Risk stratification and feature importance analysis of the multiclass HGB model. **(A)** Distribution of samples across risk categories by phenotype, with sample counts shown within each bar segment. **(B)** Top 20 taxa ranked by feature importance in the Random Forest model.

Feature importance analysis revealed that specific microbial taxa, including *Fusobacterium nucleatum* and various *Bacteroides* species, were among the most informative features for classification. Age and smoking status also emerged as important predictive features (**Figure 6B**).

## Discussion/Conclusion

Our platform is unique since it provides curated metadata, pre-computed results of publicly available samples, an analysis pipeline for analysing customer samples, cohort building, data visualization, and user-friendly interface at the same time, while none of the other microbiome databases on the market provide all these features. For example, GMRepo is a well-curated human gut microbiome database with precomputed results; however, it does not provide an on-platform analysis pipeline nor allows for sample submission and cohort building (31,32). Qiita, an open-source microbial study management platform, does provide most of the features we provide; however, it is difficult to navigate, and an average user would likely need to read through a lengthy tutorial before they can successfully use this platform (27). NMDC is a well-maintained platform that integrates and standardizes multi-omics microbiome data; however, its cohort building feature is limited as it focuses mostly on environmental samples and only includes a limited set of metadata (61).

We conducted three case studies to benchmark results achieved using our platform. In case study 1, we used Orion to analyse a colorectal cancer (CRC) dataset and compared it to the results reported in the original study (55). We observed more unique species with the Orion analysis and partial concordance in differentially abundant species between CRC and healthy groups with the original study, likely reflecting the fact that the original study employed different analysis tools and reference databases in 2019 (55).

Secondly, the Parkinson’s Disease (PD) case study demonstrates the value of the Orion Microbiome Database as a harmonized platform for investigating complex disease-associated microbiome patterns, using PD as an example. By integrating data from 742 fecal microbiome samples spanning three BioProjects, Orion addressed challenges that typically impede secondary analysis of public datasets, namely fragmented repositories, inconsistent metadata reporting, heterogeneous sequence analysis pipelines, and limited cross-study comparability. The ability to perform standardized, reproducible analyses across cohorts underscores Orion’s potential as a unifying resource for large-scale microbiome investigations.

A first objective of this analysis was to examine how sequencing platform and study-specific factors might influence microbial community profiles in PD. Our results highlight notable taxonomic differences across projects, pointing to the pervasive impact of technical variation. For instance, samples sequenced on the Illumina NovaSeq 6000 (PRJNA834801) showed higher relative abundances of *C. cayetanensis*, *B. wexlerae*, and *B. faecis*, while *Alistipes putredinis* was significantly reduced. Conversely, the HiSeq X Ten cohort (PRJNA433459) was enriched in *Bacteroides* species (*B. thetaiotamicron, B. ovatus, B. uniformis*) and *P. dorei*, but depleted in *A. muciniphila*, *C. cayetanensis*, *F. saccharivorans*, and several *Blautia* species. Such contrasts underscore a well-documented issue in microbiome science: sequencing technology, library preparation, and project-specific methodologies can introduce systematic biases that rival or even exceed biological signals (62). Without harmonization, these sources of variation complicate cross-study comparisons and reduce reproducibility, highlighting the importance of platforms like Orion that standardize downstream analyses with post hoc computational correction to adjust for batch effects (63).

In addition to technical variation, Orion enabled comparisons of microbiome composition between PD and healthy phenotypes across aggregated datasets. Several taxa emerged as consistently associated with PD. Increased abundances of *B. wexlerae*, *C. cayetanensis*, *Clostridiales bacterium*, *uncultured Oscillospiraceae* and *Eubacteriales* taxa, and *F. saccharivorans* were observed, alongside decreased abundances of *B. thetaiotamicron* and *Phocaeicola massiliensis* (**Figure 3**). These findings align with emerging literature suggesting that PD is associated with disruption of gut microbial communities, particularly taxa involved in short-chain fatty acid (SCFA) production, mucin degradation, and immune modulation (64–67). For example, *Blautia* species are known butyrate producers, and their increased abundance in PD may reflect compensatory changes in microbial metabolism (68). Similarly, the depletion of *B. thetaiotamicron*, a key glycan degrader, has been reported in prior PD studies and may contribute to altered gut barrier function and host-microbe interactions (69). Together, these results highlight Orion’s capacity to replicate known associations.

The dual findings from this case study emphasize two broader implications for microbiome research. First, sequencing technology and methodological heterogeneity remain critical confounders in cross-cohort analyses. Differences observed between Illumina platforms illustrate how technical variation can obscure or mimic biological effects if not carefully accounted for. Second, despite these challenges, standardized pipelines and harmonized metadata, as provided by Orion, enable researchers to extract biologically meaningful insights that extend beyond the scope of individual studies. By systematically disentangling technical artifacts from genuine microbial associations, Orion facilitates robust meta-analyses and accelerates discovery of disease-related microbial signatures.

In the context of PD, these results reinforce the emerging view that the gut microbiome plays a role in disease pathophysiology, potentially through pathways involving SCFA metabolism, mucosal integrity, and immune signalling (70,71). At the same time, the platform-specific differences observed underscore the caution needed when interpreting microbiome-disease associations from isolated datasets. Ultimately, the integration and harmonization enabled by Orion can help overcome these barriers, supporting more reproducible and generalizable insights into host-microbe interactions. As more datasets are incorporated and analytical tools expanded, Orion has the potential to serve as a cornerstone for reproducible, large-scale microbiome research in PD and other complex diseases.

Finally, our CRC prediction model case study further demonstrates Orion’s ability to generate clinically meaningful insights from curated microbiome cohorts. Using Orion’s cohort-building functionality, we assembled a harmonized dataset of 2,392 samples spanning healthy controls, adenoma cases, and CRC patients from multiple geographically diverse studies. This integration, facilitated by Orion’s standardized metadata framework and unified taxonomic profiling, enabled construction of both binary and multiclass prediction models without the technical heterogeneity challenges typically encountered when aggregating public microbiome data. This capability directly addresses a critical barrier in microbiome-based diagnostic development: the ability to rapidly compile large, well-annotated cohorts from fragmented repositories.

Our binary classifier achieved strong discrimination between CRC and healthy states, with AUROC values exceeding 0.94 across cross-validation and held-out test evaluation (**Table 4**). The multiclass model, which incorporated the adenoma stage, effectively separated all three phenotypes with strong discriminative performance (AUROCs >0.91) and achieved exceptional adenoma detection (94.1% recall, **Table 5**). This latter result is especially noteworthy, as adenoma detection represents a critical intervention point in the adenoma-carcinoma sequence yet remains a major unmet clinical need (72), with conventional non-invasive screening methods such as fecal immunochemical testing (FIT) typically achieving lower sensitivities for precancerous lesions (73). By explicitly modeling the adenoma stage, our multiclass framework addresses this gap in CRC screening.

The risk stratification framework embedded in our models demonstrates translational potential for clinical implementation (**Figure 6A**). The High Risk category, predominantly composed of CRC patients (69.9%), captured 66.4% of all CRC cases, while the Medium Risk tier consisted primarily of healthy individuals (87.5%). This stratification could serve as a triage tool to prioritize colonoscopy referrals for high-risk individuals, thereby improving the cost-effectiveness and patient acceptance of CRC screening programs (74).

The taxonomic features driving model predictions corroborate established literature on microbiome involvement in colorectal carcinogenesis (**Figure 6B**). *Fusobacterium nucleatum* and *Parvimonas micra*, both consistently associated with CRC progression, emerged as key predictive taxa: *F. nucleatum* is enriched in adenoma and carcinoma tissues and is associated with microsatellite instability, low T-cell infiltration and poorer survival; it also expresses the adhesin FadA, enabling invasion of colonic epithelial cells and activation of Wnt/β-catenin signaling (75). In experimental models P. micra promotes colonocyte proliferation and Th17-mediated immune responses and is associated with poorer survival in CRC patients (76). Conversely, *Anaerostipes hadrus* and *Blautia* species emerged as protective markers: butyrate-producing commensals such as *Anaerostipes* utilize lactate and acetate to generate butyrate, and butyrate has been shown to maintain gut barrier integrity, limit pro-inflammatory cytokines and inhibit oncogenic pathways including Wnt signaling (77); the gut commensal *Blautia* stimulates colonic mucus growth through production of propionate and acetate, supporting barrier function (78). These observations support the driver-passenger model of CRC, in which early “driver” bacteria induce DNA damage and inflammation, while opportunistic “passenger” bacteria, including *F. nucleatum* and *P. micra*, colonize tumors and further promote progression (75).

Beyond microbial features, age and smoking status emerged as important predictive features in our models (**Figure 6B**). This finding aligns with well-established epidemiological evidence identifying both factors as major CRC risk determinants: incidence rises sharply after age 50, and smoking increases risk through carcinogenic compound exposure and chronic inflammation (79). The integration of these clinical risk factors alongside microbiome signatures demonstrates that microbiome-based classifiers can capture both host demographic risk and microbial dysbiosis patterns associated with CRC progression.

Critically, this case study shows Orion’s unique value proposition: the platform’s curated metadata, standardized processing pipeline, and cohort-building interface enabled rapid assembly and analysis of a multi-study dataset that would otherwise require extensive manual curation and harmonization. The ability to seamlessly integrate samples across BioProjects, apply consistent quality filters (age ≥30 years, sequencing depth >1M reads), and exclude non-biological confounders through Orion’s interface accelerated translational research and ensured reproducibility.

Currently, only human gut samples are available on Orion; however, we are planning on expanding our database to first include other human samples, such as skin, oral, vaginal, etc., and eventually to include other animal samples and environmental samples as well. In the long run, we aim to server the entire microbiome research community with different interests and needs as a centralized place to retrieve data and conduct various analysis.

---

**Supplementary Table 1.** Summary of public cohorts.

| Name | Description | Samples | Filters | Publisher |
| --- | --- | --- | --- | --- |
| Active Pregnancy | Stool samples from actively pregnant women with variance in pregnancy term. | 70 | Phenotype = Pregnant | Zymo Research |
| Kidney Failure | Stool samples from chronic kidney failure patients | 223 | Phenotype = Kidney Failure; Chronic | Zymo Research |
| Parkinson's Disease | Fecal and gut samples from patients with a confirmed Parkinson's diagnosis. Samples were collected at various states of disease progression. | 488 | Phenotype = Parkinson's Disease | Zymo Research |
| Colorectal Cancer | Fecal samples from patients with colorectal cancer. | 660 | Phenotype = Colorectal Cancer | Zymo Research |
| Autistic Spectrum | Stool samples from patients diagnosed on the autism spectrum from patients under 10 years old. | 323 | Phenotype = Autistic Spectrum Disorder, Age < 10 | Zymo Research |
| Healthy Infants | Fecal samples from healthy infants. Healthy is defined as no other associated phenotypes and healthy infantile weights. Infant ages range from newborn to 2 years of age. | 36 | Phenotype = Healthy, Age <= 2 | Zymo Research |
| Liver Cirrhosis | Fecal samples from patients with confirmed liver cirrhosis with varying degrees of severity and duration. | 144 | Phenotype = Liver Cirrhosis | Zymo Research |
| Crohn's Disease | Fecal and gut samples from Crohn's disease patients with a varying degree of disease activity. | 303 | Phenotype = Crohn's Disease | Zymo Research |
| Healthy Adults | Fecal and gut samples from healthy adult samples. Healthy is defined as the phenotype healthy and having a BMI less than 30 to exclude clinically obese samples and excluding any co-existing phenotypes from inactive disease states. Adult is defined as any individual 18 years of age or older. | 2396 | Phenotype = Healthy, BMI < 30, Age >= 18 | Zymo Research |
| Obesity | Stool samples from obese individuals classified by a BMI greater than 30. Includes all phenotypes. | 1204 | BMI > 30 | Zymo Research |
| Rheumatoid Arthritis | Fecal and gut samples from patients with diagnosed rheumatoid arthritis | 237 | Phenotype = Rheumatoid Arthritis | Zymo Research |
| Ulcerative Colitis | Fecal samples collected from Ulcerative Colitis patients. | 257 | Phenotype = Ulcerative Colitis | Zymo Research |

